# Validating Pore Size Estimates in a Complex Microfibre Environment on a Human MRI System

**DOI:** 10.1101/2021.03.27.437304

**Authors:** Chu-Chung Huang, Chih-Chin Heather Hsu, Feng-Lei Zhou, Slawomir Kusmia, Mark Drakesmith, Geoff J.M. Parker, Ching-Po Lin, Derek K. Jones

**Affiliations:** Institute of Cognitive Neuroscience, School of Psychology and Cognitive Science, East China Normal University, Shanghai, China; Center for Geriatrics and Gerontology, Taipei Veterans General Hospital, Taipei 11217, Taiwan; Institute of Neuroscience, National Yang Ming Chiao Tung University, Taipei 11221, Taiwan; Centre for Medical Image Computing, Department of Computer Science, University College London, London, WC1V 6LJ, UK; School of Pharmacy, University College London, London WC1N 1AX, UK; Cardiff University Brain Research Imaging Centre (CUBRIC), Cardiff University, Cardiff, United Kingdom; Centre for Medical Image Computing (CMIC), University College London, London, United Kingdom; Epilepsy Society MRI Unit, Chalfont St Peter, United Kingdom; Department of Neuroinflammation, Queen Square Institute of Neurology, University College London, London, WC1V 6LJ, UK; Bioxydyn Limited, Manchester, UK

**Keywords:** Diffusion MRI, Microstructure, Phantom, Electron Microscopy, Crossing Fibre, Diameter

## Abstract

**Purpose:** Recent advances in diffusion-weighted MRI provide ‘restricted diffusion signal fraction’ and restricting pore size estimates. Materials based on co-electrospun oriented hollow cylinders have been introduced to provide validation for such methods. This study extends this work, exploring accuracy and repeatability using an extended acquisition on a 300 mT/m gradient human MRI scanner, in substrates closely mimicking tissue, i.e., non-circular cross-sections, intra-voxel fibre crossing, intra-voxel *distributions* of pore-sizes and smaller pore-sizes overall.

**Methods:** In a single-blind experiment, diffusion-weighted data were collected from a biomimetic phantom on a 3T Connectom system using multiple gradient directions/diffusion times. Repeated scans established short-term and long-term repeatability. The total scan time (54 minutes) matched similar protocols used in human studies. The number of distinct fibre populations was estimated using spherical deconvolution, and median pore size estimated through the combination of CHARMED and AxCaliber3D framework. Diffusion-based estimates were compared with measurements derived from scanning electron microscopy.

**Results:** The phantom contained substrates with different orientations, fibre configurations and pore size distributions. Irrespective of one or two populations within the voxel, the pore-size estimates (~5μm) and orientation-estimates showed excellent agreement with the median values of pore-size derived from scanning electron microscope and phantom configuration. Measurement repeatability depended on substrate complexity, with lower values seen in samples containing crossing-fibres. Sample-level repeatability was found to be good.

**Conclusion:** While no phantom mimics tissue completely, this study takes a step closer to validating diffusion microstructure measurements for use *in vivo* by demonstrating the ability to quantify microgeometry in relatively complex configurations.

## Introduction

Obtaining reliable *quantitative* tissue microstructure information using non-invasive magnetic resonance imaging has long been a holy grail in microstructural MRI^1,2^. Improvements in gradient hardware^3,4^ give increased sensitivity to smaller water displacements, and higher SNR per unit time, while improvements in modelling^2^ can potentially yield higher specificity to compartment-specific properties. Two measures of particular interest are: (i) the fraction of the signal that comes from spins trapped within pores, known as the ‘restricted signal fraction’^5^; and (ii) the inner-diameter (pore-size) of restricting geometries^6,7^. In white matter, for example, the former is taken as an indicator of ‘axon density’, while the latter is taken as an indicator of axon diameter - one of the major factors influencing the speed of action potentials^8,9^.

Most previous validations of such measurements have estimated diameters in tightly controlled architectures (e.g., using a phantom comprising synthetic fibres all with the same orientation, or in the mid-line of the corpus callosum where the fibres are largely co-aligned) on small-bore preclinical scanners (exploiting the strong gradients that typically accompany such systems)^7,10–18^. In comparison, there is much less work validating measurements in more complex substrates using MRI systems designed for clinical use, which is essential for pushing forward *in vivo* microstructural quantification in human tissue.

*Ex vivo* / post-mortem brain samples might be the most direct route for validation since, by definition, they reflect the real physical complexity of the tissue, albeit with limitations imposed by changes in relaxation times, diffusivities, and tissue shrinkage^19^. However, lack of parametric control over tissue properties, such as size, shape or distribution makes the systematic study of the performance characteristics of a microstructural pipeline more challenging.

The ability to <u>specify</u> the microstructural properties of a substrate *a priori* can potentially facilitate the design of far more efficient experiments to ascertain accuracy and precision. Numerical / *in silico* phantoms have been used to simulate different substrates by modelling diffusion properties with different geometries^20,21^. However, digital phantoms are generally over-simplistic in several respects, including the fact that they do not simulate acquisition conditions faithfully^22^. To the best of our knowledge, this can only be achieved feasibly through actual scanning of physical phantoms comprising synthetic substrates^10,23–25^. As discussed by Fan, et al.,^10^ while physical phantoms can never fully replace *ex vivo* samples in reflecting real tissue status, they serve as an important step between *in silico* simulations and actual construction of biological tissues.

Using a hollow textile filament (or ‘taxon’) phantom, Fan, et al.,^10^ validated non-invasive pore size estimates on a human MRI system with ultra-strong gradients (up to 300 mT/m). Sampling over a wide range of diffusion times and gradient strengths, they estimated inner diameter and restricted signal fraction using a simplification of the *AxCaliber* model^7^ that, as with *ActiveAx*^6^, fits for a single pore diameter. Their results showed good agreement with the known phantom properties, supporting the feasibility of estimating microfibre pore size on a clinical MRI system. However, the data-acquisition took 38 hours, and the phantom comprised fibres with a: (i) single orientation; (ii) circular cross-section; and (iii) a single, relatively large (compared with diameters typically found in the brain^26^), diameter of 12 μm. While this work therefore represents an important step in understanding the capabilities of emerging hardware and modelling frameworks to recover microstructural parameters on a clinical system, it is important to extend the validation framework into more complex substrates, moving towards architectures that mimic tissue properties *in vivo*. Moreover, for full clinical translation, exploring the fidelity of microstructural estimates with shorter acquisition protocols is essential.

To approach the kinds of microstructural geometries found *in vivo*, and to achieve the parametric control of properties such as pore size, shape, density and orientation, Zhou, et al.,^27^ developed the manufacture of co-electrospun microfibres from highly hydrophilic hollow polycaprolactone (PCL). Critically, this approach has a stochastic element, thereby introducing a *distribution* in the cross-sectional shape of individual pores, and facilitates control over pore size and orientation^25^. This approach has been used to create ‘axon-mimicking’ fibres^25,27^ that resulted in anisotropic diffusion of water within them. Moreover, by tuning the fibre inner-diameter, the authors previously demonstrated control over diffusion tensor MRI-based properties such as fractional anisotropy and radial diffusivity^25^ and used these materials to help characterise signals from multidimensional diffusion encoding acquisitions^28^.

Again, while work such as described above represent steps towards validating estimates of pore size, several caveats remain. In previous validations of *AxCaliber*, the free diffusivities of the liquid and inner pore diameter were known *a priori*. Moreover, measurements were limited to samples with a single and known orientation (physical phantom/corpus callosum), limiting the generalizability of the validation of quantitative measurements across a whole organ such as the brain. It is also important to ensure that such measurements have excellent short-term and long-term repeatability, to ensure that any observed changes in the signal/measurement reflect true changes in the substrate being imaged, rather than a perturbation introduced through noise/scanner instability/instabilities in the data-processing pipeline. To facilitate this, it is important to have a substrate that will not change its physical properties over time, but which also mimics the physical properties of the target substrate of interest (e.g., white matter).

To address these issues, we aimed to validate *AxCaliber* estimations of microstructural parameters using co-electrospun substrates containing microfibres with <u>unknown</u> (to a subset of the authors) distributions of size, shape and orientation and complexity (i.e., number of distinct compartment populations). This study was completed in a single-blind manner to prevent any potential bias in estimates, pre-processing or *post hoc* inference. Thus, an essential feature of this study was that any information on the phantom microstructure was totally withheld from a subset of the authors (C-CH, C-CHH, SK, DKJ) until all data acquisition, analysis and final estimations were complete. To maintain translational relevance to *in vivo* applications, we used a diffusion-weighted imaging protocol with a total scan time less than one hour (54 mins). Constrained spherical deconvolution (CSD)^29^ was used to estimate the number of distinct fibre orientations and a single-parameter continuous Poisson distribution model^30^ within the Axcaliber3D^31^ framework to fit the range of pore sizes in the biomimetic phantom.

## Method

### Experimental Design

At the beginning of this single-blind validation study, a subset of the authors knew only that the phantom contained 6 tubes with hollow microfibres and 1 tube with pure liquid medium produced according to the method of Zhou, et al.,^27,32,33^. Other than a numbering system to reference each tube, it was impossible to differentiate between the samples with the naked eye. The construction of the phantom is shown in Figure 1A, and ‘sagittal’ and ‘axial’ sections of the phantom through a diffusion-weighted MRI (dMRI) are shown in Figure 1B. Figure 1C shows a schematic overview of the experimental design, and full protocol details are described in the following sections.

**Figure 1.**
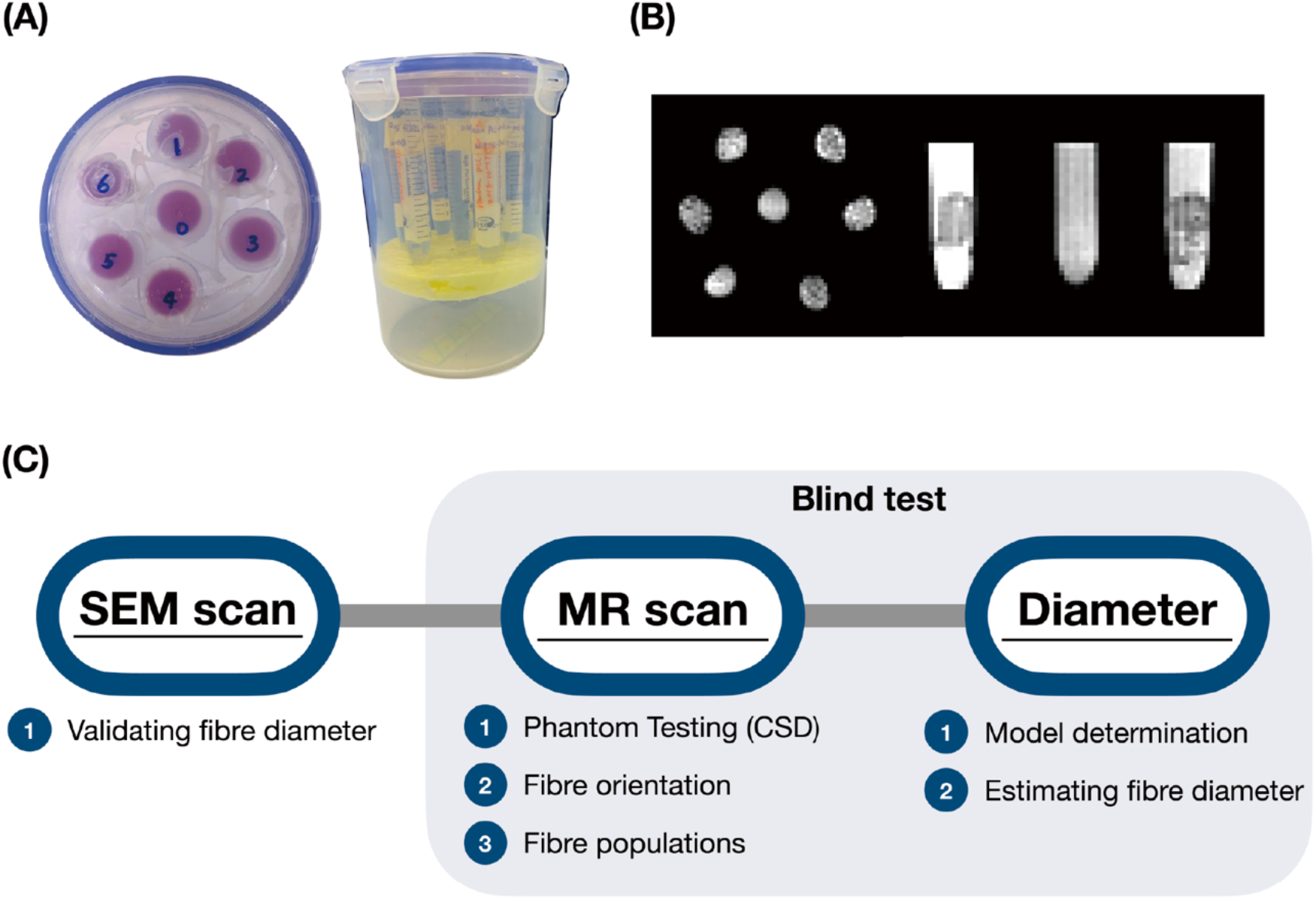
The overview of the diffusion phantom and the design of the single-blind experiment. (A) There were seven tubes in the phantom container, each tube held one or two block-samples of fibre phantom with liquid filled (the characteristic of filling medium was unknown during experiments). (B) The sagittal and coronal slices of the diffusion-weighted image of the phantom are shown to demonstrate the voxels with anisotropic phantoms. (C) The flowchart describes the design of the single-blind experiment.

### Phantom construction and fibre characterization

The co-electrospinning setup used to generate the phantom has been detailed previously^34^. Briefly, a coaxial spinneret with two concentric needles was filled with a solution of 10 wt.% polycaprolactone (PCL, Mn = 80k g/mol) with 1 wt. % polysiloxane-based surfactant (PSi) in CHCl_3_/DMF (8/2 w/w) (outer needle, inner diameter = 1.19 mm) and 4 wt.% polyethylene oxide (PEO, Mv = 900k g/mol) in water (inner needle, inner diameter = 0.41 mm)^27^. The outer needle was connected to the positive electrode of a DC voltage power supply, while the fibre collector was grounded. A voltage of 9 or 15 kV was applied to generate uniaxially or randomly aligned fibres respectively. A rotating drum (diameter = 11 cm), spinning at 800 or 10 revolutions per minute, was placed at a distance of ~5 or 14 cm from the tip of the concentric needles as a collector. To allow the fibre deposition to be spread uniformly, the collector was positioned on a translational *x-y* stage, moving left and right at 1 mm/s. The flow rates of the outer and inner solutions were fixed at 3 and 1 mL/h, respectively. The translation distance of the *x-y* stage was 30 or 55 mm for uniaxially or randomly aligned fibres.

Six phantom samples labeled 1-6 were constructed by packing one (Tubes #1, #5 and #6) or two blocks (Tubes #2, #3 and #4) comprising ~24 fibre layers (10 mm × 10 mm) into a 15 ml centrifuge tube filled with deionized water. The same co-electrospun substrate was used for the construction of Tubes #1 and #6, where the fibres comprising the block were stacked in an interleaved fashion, crossing at 45° (Tube #1) and 90° (Tube #6); fibres comprising the blocks in Tube #2 and #3 were randomly oriented; fibres *within* each block inside Tubes #4 and #5 were uniaxially aligned (0°), but in Tube #4, the two distinct blocks (made from the same substrate) were oriented at 90° to each other. Six phantom samples were initially placed into a vacuum-degassing chamber to remove air bubbles before they were assembled into a cylindrical plastic container (inner diameter: ~140 mm; height: ~180 mm); The six tubes (#1-6) were spaced equally around the circumference of the container and one tube containing only deionized water (labeled 0) was placed at the centre. Due to the single-blind study design, the specification/configuration of the phantom was not revealed until the MRI acquisition and data analyses were complete.

### MRI experiment

The diffusion-weighted phantom scans were performed on the 3T Connectom MRI system (maximum gradient strength = 300 mT/m) using a Siemens 32-channel head coil at the Cardiff University Brain Research Imaging Centre (CUBRIC). The phantom container was placed along the scanner’s y-axis (vertical orientation and perpendicularly to the static magnetic B0 field), so that any air bubbles floated to the top of the tubes, and was immobilized with cushions to minimize vibrations during scanning.

The same imaging protocol was applied four times to evaluate the repeatability of diameter estimates. The first two scans (Scan 1 and Scan 2) were conducted on 20^th^ May 2019 and 3^rd^ June 2019, whereas the other two scans (Scan 3a and Scan 3b) were a pair of immediate scan-rescan acquisitions conducted on 16^th^ June 2020. All scans were performed under ambient conditions and the temperature was not recorded. The protocol comprised two diffusion frameworks, the Composite Hindered And Restricted ModEl of Diffusion (CHARMED)^5^ and the *AxCaliber3D* framework^31^. The CHARMED model considered the diffusion signal to arise from a combination of hindered and restricted diffusion components and fitted the data to the composite model, with a fixed diameter distribution of fibre to estimate signal fractions, diffusivity parameters and axonal orientations. In contrast, the *AxCaliber* model expands CHARMED by introducing the diameter distribution of restricted cylindrical fibres as an unknown function to estimate but, in its original implementation, only considers diffusion-encoding along a single axis, assumed to be orthogonal to the fibre orientation. By combining CHARMED and AxCaliber, *AxCaliber3D* enables axon diameter distributions to be recovered for more complicated fibre configurations, and with arbitrary orientation of the fibres with respect to the diffusion encoding. Both datasets were acquired using a diffusion-weighted spin-echo blipped-CAIPI (EPI) sequence^35^ with 1.5 mm isotropic resolution, with parameters summarized in Table 1. For the CHARMED acquisition, the diffusion gradient pulse duration and the diffusion time were both fixed, and the gradient amplitude was varied between 51 mT/m and 281 mT/m resulting in b-values ranging from 200 to 6000 s/mm^2^. In each shell, the diffusion-encoding gradient directions were uniformly distributed over the unit sphere according to Jones, et al.,^36^. For AxCaliber3D, images with six different diffusion times were acquired using a fixed gradient pulse duration and varying the prescribed b-value between 2200 and 25500 s/mm^2^, with a maximal gradient amplitude of 288 mT/m. In each b-value shell, data were acquired over 30 uniformly-distributed encoding directions. A total of 24 b = 0 s/mm^2^ images were interleaved between the different b-shells to allow for the correction of signal drift. The total acquisition time was 54 min. Other imaging parameters included a field-of-view of 128 mm x 128 mm, 30 continuous slices, with an isotropic voxel size of 1.5 mm, simultaneous multi-slice factor of 2, partial Fourier of 6/8, and no GRAPPA was applied.

**Table 1.** Diffusion-weighted image protocol.

| CHARMED (TR/TE = 4500/74ms) |  |  |  |  |  |
| --- | --- | --- | --- | --- | --- |
| b-values<br>(s/mm<sup>2</sup>) | # of<br>diffusion<br>directions | Gradient<br>Strength<br>(mT/m) | Range of q-<br>value (μm<sup>-1</sup>) | Gradient pulse<br>duration (δ, ms) | Diffusion<br>times (Δ, ms) |
| 200 | 20 | 51.30 | 0.015 | 7 | 24 |
| 500 | 20 | 81.12 | 0.024 |  |  |
| 1200 | 30 | 125.7 | 0.038 |  |  |
| 2400 | 61 | 177.7 | 0.053 |  |  |
| 4000 | 61 | 229.4 | 0.068 |  |  |
| 6000 | 61 | 281.0 | 0.084 |  |  |
| 3D-AxCaliber (TR/TE = 5000/138ms) |  |  |  |  |  |
| 2200/4400 | 30 for<br>each<br>shell | 200.1/283.0 | 0.060/0.084 | 7 | 18 |
| 3600/7150 |  | 204.0/287.5 | 0.061/0.086 |  | 27 |
| 4900/9800 |  | 203.7/288.1 | 0.061/0.086 |  | 36 |
| 6200/12400 |  | 203.5/287.9 | 0.061/0.086 |  | 45 |
| 8400/16750 |  | 203.8/287.9 | 0.061/0.086 |  | 60 |
| 12750/25500 |  | 203.6/288.0 | 0.061/0.086 |  | 90 |

### Data pre-processing and analysis

The regions of interest (ROIs) used for model fitting were selected manually from the cross-sectional images of the tubes through the following steps: 1) thresholding the b = 0 s/mm^2^ images with intensity higher than 10% of its maximal signal intensity to avoid processing background noise; 2) separating the thresholded binary mask image spatially into 9 ROI components for each fibre sample in the tubes (labelled as Tube #1, #2a, #2b, #3a, #3b, #4a, #4b, #5, #6, corresponding to the label on the phantom tubes as shown in Figure 1A); 3) cropping the ROI along the axis parallel to the tube orientation to ensure that it only covered the anisotropic samples in the tube; 4) Eroding the ROIs by two voxels to eliminate inhomogeneous partial-volume voxels at the boundaries between the phantom material and the plastic tube containing the material.

For both the CHARMED and AxCaliber3D datasets, the signal pre-processing involved: 1) Denoising^37^; 2) Drift correction^38^; 3) Eddy current distortion^39^; 4) Gradient nonlinearity distortion^40^; and 5) Correction for Gibbs-ringing artifacts^41^.

The b = 2400 s/mm^2^ (61 directions) shell data of the pre-processed CHARMED data were used to derive the fibre orientation distribution function (fODF) via CSD with l_max_ = 8 in MRtrix3 (http://www.mrtrix.org/). The number of unique fODF peaks in each sample was extracted via Newton optimization^42^. The threshold was set to 0.1 absolute amplitude of the fODF and above 33% of the maximum amplitude of each voxel to exclude small peaks^43^.

The AxCaliber3D framework^31^, embedded into the Microstructure Diffusion Toolbox (MDT, https://github.com/robbert-harms/MDT), was used to estimate the microfibre inner diameters. MDT includes a model-cascade approach^44^ that shortens the overall run time and improves fitting accuracy and precision. Initially, CHARMED data were used to model the signal using one hindered diffusion compartment (using a zeppelin diffusion tensor) and one or two restricted diffusion compartments (based on the estimated number of fibre populations from the CSD analysis) using van Gelderen’s^45^ expression for restricted diffusion in a cylinder. The estimated fibre peak orientations then served as prior fixed parameters and initial starting estimates of the restricted diffusion signal fractions for the fitting of the AxCaliber3D model. The total measured signal decay was assumed to be a sum of diffusion-weighted signal decays for each pore size weighted by their respective area-weighted probability and the pore-size populations were modelled with a continuous Poisson diameter distribution^30^, yielding an average pore-size for each voxel in the ROI.

### Scanning electron microscopy of the phantom

The surface morphology and cross-sections of co-electrospun fibres were observed using a FEI Quanta 650 field emission gun scanning electron microscope (SEM) with an accelerating voltage of 5 kV. The co-electrospun fibre specimens were coated with a gold-palladium film to increase their conductivity and the fibre strips were cut using a sharp scalpel in liquid nitrogen for imaging their cross-sections. ImageJ (<u>imagej.nih.gov/ij</u>) was used to measure the pore size (fibre inner diameters) using its “Pore Measurement” function. For each sample, pore sizes (areas) were automatically measured from 5 different SEM images and manually converted into the fibre inner diameters within the sample with the assumption of circular pores^33^. The area-weighted fibre inner diameters and fractions were calculated using a method reported previously^46^. Those responsible for the phantom manufacture (FZ, GP) noted that, because fibre deposition could not be controlled precisely during the co-electrospinning process, some large ‘extra-fibre’ pores were formed randomly and frequently in the phantom (see Figure 2 and Supplementary Information Figure S1). In these spaces, for the diffusion times used here, the spins would not experience hindrance/restriction during their displacement, effectively leading to a third mode of diffusion, i.e., ‘free water’. Estimating all pore sizes, irrespective of dimension (and including these larger pores) would give a false impression of restricting pore size, and so the estimated restricting fibre volume fractions from SEM were estimated by dividing the total area of pores with diameter less than 15 μm, by the total area of all pores (i.e., pores with diameters in the range: [0, ∞]), – see Table 2).

**Figure 2.**
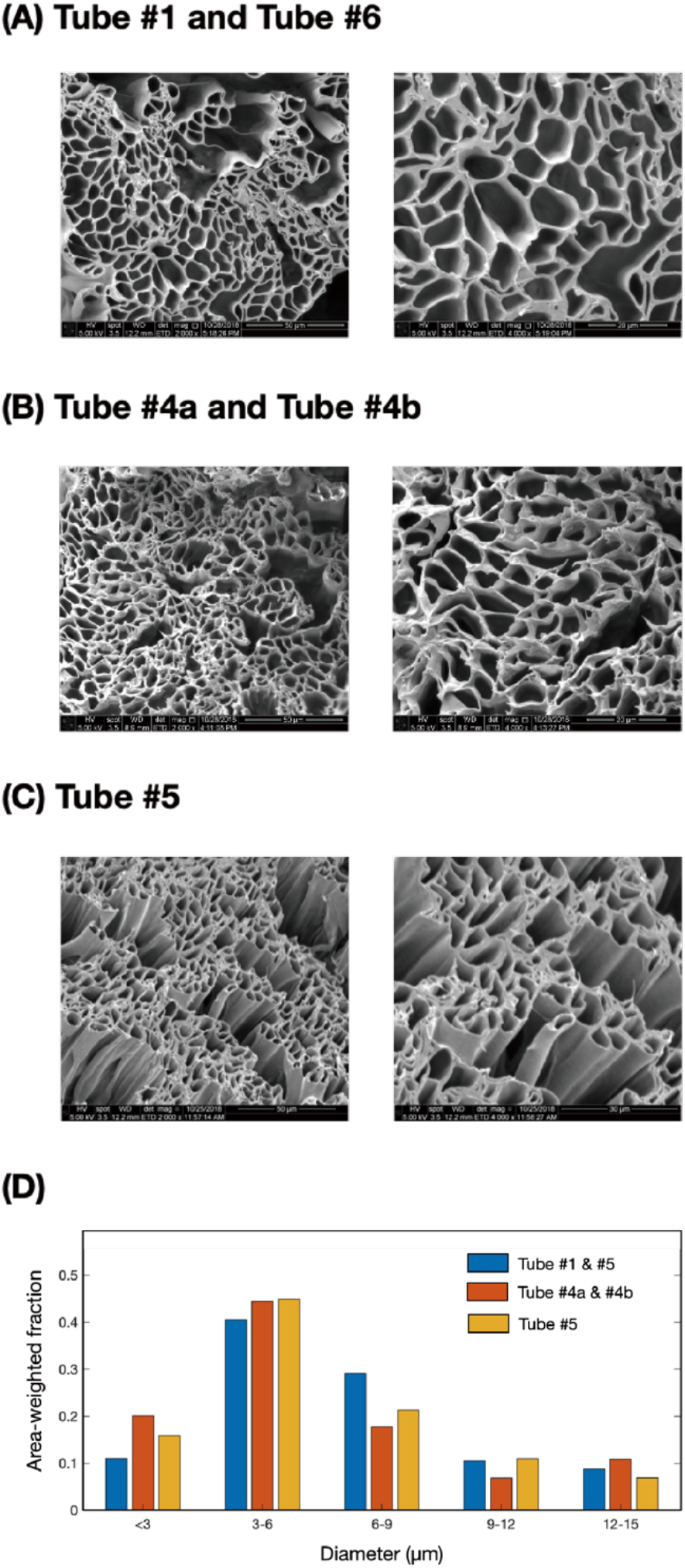
SEM images of Co-Electrospun PCL-Psi fibre phantom. (A-C) show the SEM images with low (left) and higher (right) resolution. In (D), the area-weighted fractions of each sample pore size are shown in blue (Tube #1 and Tube #6), orange (Tube #4a and #4b), and yellow (Tube #5).

**Table 2.** phantom properties estimated by dMRI and SEM.

|  | Scan 1<br>(May 20<sup>th</sup> 2019) |  | Scan 2<br>(June 3<sup>rd</sup> 2019) |  | SEM |
| --- | --- | --- | --- | --- | --- |
|  | P1 | P2 | P1 | P2 |  |
| **Tube #1** |  |  |  |  |  |
| Pore Size (μm) | 4.97 (4.15, 5.62) | 5.19 (4.45, 5.72) | 5.54 (4.03, 7.19) | 6.02 (4.34, 7.95) | 6.01 (4.42, 8.03) |
| Signal/Volume Fractions<sup>a</sup> | 0.17 (0.15, 0.20) | 0.17 (0.15, 0.19) | 0.18 (0.15, 0.22) | 0.17 (0.15, 0.19) | 0.36 (0.34, 0.62) |
| Angle (°) | 50.4 (47.6, 54.8) |  | 48.4 (45.7, 51.6) |  | 45<sup>b</sup> |
| **Tube #6** |  |  |  |  |  |
| Pore Size (μm) | 4.84 (4.17, 5.57) | 4.72 (3.94, 5.47) | 5.24 (4.36, 6.72) | 4.90 (4.09, 6.43) | 6.01 (4.42, 8.03) |
| Signal/Volume Fractions<sup>a</sup> | 0.17 (0.14, 0.19) | 0.15 (0.13, 0.17) | 0.17 (0.15, 0.19) | 0.16 (0.14, 0.18) | 0.36 (0.34, 0.62) |
| Angle (°) | 87.4 (84.9, 88.9) |  | 87.1 (84.4, 88.9) |  | 90<sup>b</sup> |
|  | P1 |  | P1 |  |  |
| **Tube #4a** |  |  |  |  |  |
| Pore Size (μm) | 4.98 (4.72, 5.23) |  | 5.10 (4.59, 6.99) |  | 5.13 (3.65, 7.26) |
| Signal/Volume Fractions<sup>a</sup> | 0.32 (0.24, 0.35) |  | 0.33 (0.30, 0.36) |  | 0.47 (0.46, 0.52) |
| **Tube #4b** |  |  |  |  |  |
| Pore Size (μm) | 5.04 (4.67, 5.47) |  | 5.13 (4.29, 6.12) |  | 5.13 (3.65, 7.26) |
| Signal/Volume Fractions<sup>a</sup> | 0.32 (0.27, 0.34) |  | 0.30 (0.24, 0.32) |  | 0.47 (0.46, 0.52) |
| **Tube #5** |  |  |  |  |  |
| Pore Size (μm) | 4.67 (4.47, 4.96) |  | 4.78 (4.29, 6.81) |  | 5.38 (3.96, 7.62) |
| Signal/Volume Fractions<sup>a</sup> | 0.42 (0.35, 0.44) |  | 0.40 (0.30, 0.43) |  | 0.57 (0.55, 0.59) |

Parameter estimates for the different tubes. The median (and upper and lower quartiles) are shown. “Angle” represents the crossing angle between two fibre populations. Note Tube 1 and Tube 6 contain crossing fibres so parameters for each population (P1 and P2) are shown separately. <sup>a</sup>The estimated signal fraction from dMRI represents the fitting result of restricted diffusion signal fraction, while the volume fraction of SEM represents the ratio of the total area of pores with diameter $\leq 15 \mu\text{m}$ to the total area of pores with diameter in the range $[0, \infty]$ . <sup>b</sup> The fibres in the blocks of Tubes #1 and #6 were designed to be interleaved, crossing at 45° and 90°

### Statistical Analysis

Due to the different positioning of the phantom between Scans 1, 2 and 3, it was not possible to obtain exact spatial correspondence between the ROIs across the different scans to establish long-term repeatability. Therefore, rather than comparing estimates on a voxel-by-voxel basis, we evaluated repeatability of the pore-size estimate *distribution* using the Kolmogorov-Smirnov test (KS-test) and the Jensen-Shannon distance. To compare the distributions obtained from the dMRI scans and the ground truth from SEM, we first binned the accumulated area fraction of fibre pore size into 30 bins (bin range from 0 – 15 μm, bin width = 0.5 μm). We then calculated the median value, and the first and third quantiles (Q1 and Q3) to identify asymmetric distributions. A two-sample KS-test determined whether the two samples (SEM versus dMRI, or between the repeat-scan data) came from the same continuous distribution. The intra-class correlation coefficient (ICC), within-voxel coefficient of variation (CV), and the repeatability coefficient (RC) were calculated to evaluate the repeatability of measures at both the voxel-level and sample-level of the repeat-scan^47^.

## Results

### Phantom Examination Using dMRI

#### SNR evaluation

The SNR was calculated as the mean signal within the tubes (≥10% of maximal intensity in b = 0 s/mm^2^ image) divided by the standard deviation of the background (<10% of maximal intensity in b = 0 s/mm^2^ image) using the non-diffusion-weighted (b = 0 s/mm^2^) image. The SNRs for the four scans were: 20^th^ May 2019 = 45.3, 3^rd^ June 2019 = 44.7, 1^st^ scan on June 16^th^ 2020 = 34.2, 2^nd^ scan on June 16^th^ 2020 = 34.2.

#### Fibre population information – based on CSD model

Due to the single-blind study design, the number of distinct fibre populations in each phantom tube not known before analysis. Since the number of restricted-diffusion compartments is set *a priori* in the CHARMED model, it was therefore necessary to first estimate the number of distinct fibre populations, for which CSD was employed. Figure 3 shows the CSD-estimated fibre orientations in the axial and sagittal plane of the tubes. Four distinct phantom configurations were observed: (1) randomly oriented fibres; (2) single-oriented fibre-population; (3) two fibre-populations; (4) high diffusivity, isotropic medium (in the central tube, later revealed to be water). Both Tube #1 and Tube #6 showed an obvious crossing pattern, where the proportions of voxels that contained 2 distinct fibre populations were 0.73 and 0.88, respectively. No distinct anisotropic characteristic was observed in Tube #2 and Tube #3, in which the fODF analysis suggested that more than 80% of voxels contained more than three distinct populations in each voxel. For the purposes of our study, we assumed this was consistent with a random distribution and not amenable to analysis with the CHARMED/AxCaliber3D frameworks. Moreover, we observed that Tube #4 contained two distinct fibre substrate blocks (one on top of the other), thus we labelled these two blocks as Tube #4a and Tube #4b for further analyses and reporting.

**Figure 3.**
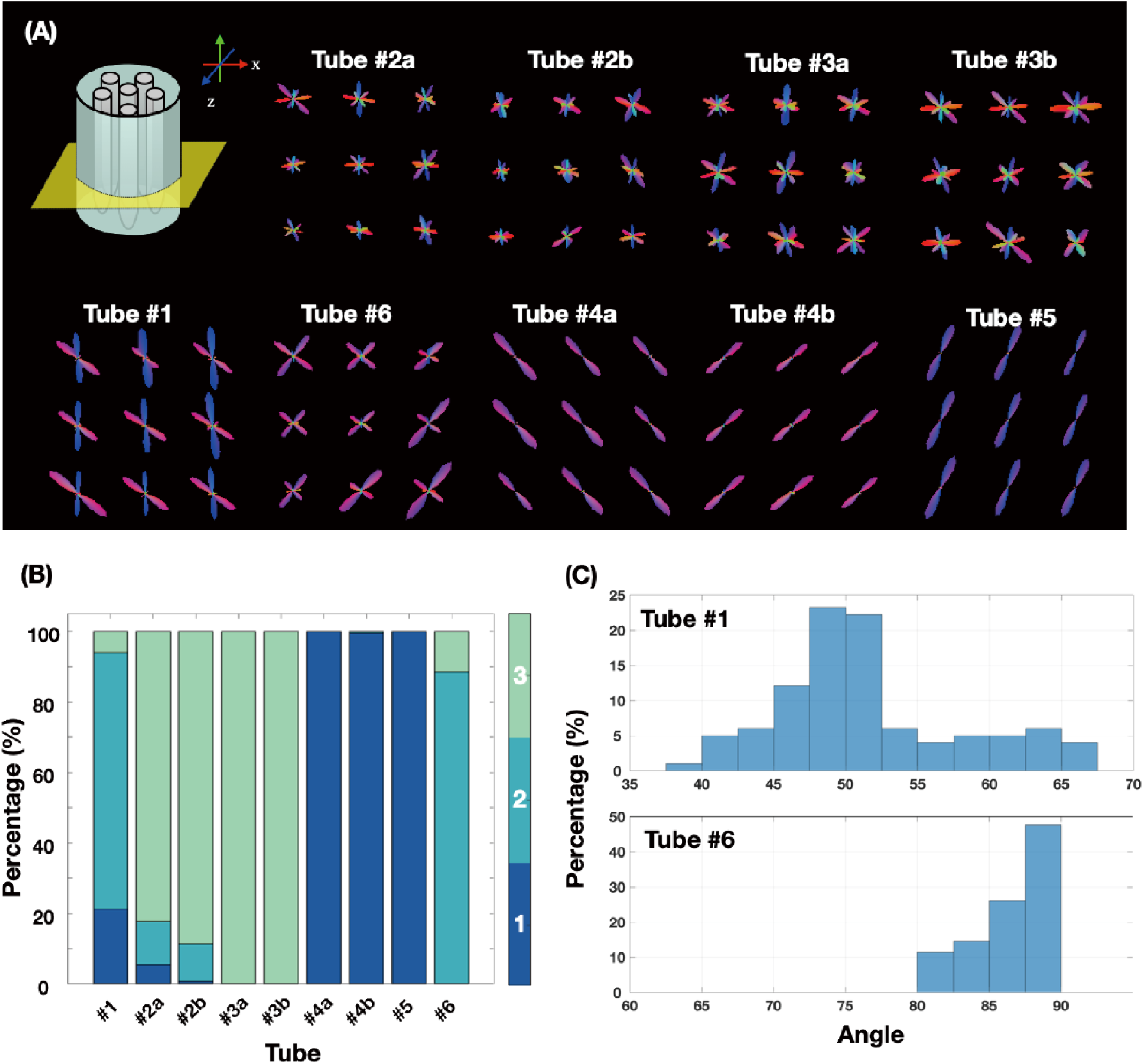
The representative measurement from the first scan on 20^th^ May 2019. Measurement of the fibre orientation density functions (fODF) in the anisotropic phantom obtained from CSD. The fibre orientations are shown with directional colour encoding in x-z plane and (B) x-y plane view. As the figure shows, Tubes #1 and #6 contain crossing fibres, Tubes #4a, #4b, #5 contain a single orientation, while Tube #2a, #2b, #3a, and #3b appear to contain randomly oriented fibres. (C) The number of unique fODF peak orientations in each sample voxel with the threshold of 0.1 absolute amplitude and 33% of the maximum amplitude. The colour represents the number of unique peaks, where blue = 1, cyan = 2, and green represent 3 or more peaks. (D) The histogram of the estimated angle between fibres in each voxel in Tubes #1 and #6 (left and right, respectively).

#### Angle information – based on CHARMED

The median crossing angles of Tube #1 and Tube #6 were estimated using the CHARMED framework to be 50.37° and 87.36° respectively (Figure 3C and Table 2). No crossing fibre configuration was observed in Tube #4a, Tube #4b or Tube #5 (Figure 3A-B).

#### Pore sizes – based on 3D-AxCaliber

Due to parameter explosion, voxels with a large number of randomly aligned fibres (e.g., Tubes #2 and #3) are not amenable to analysis by the Axcaliber3D framework. Thus, we only reported fibre diameter estimates in samples identified as containing one or two fibre orientations (Tubes #1, #4a, #4b, #5, and #6). Table 2 shows the estimated median fibre diameters and the estimated restricted signal fraction. In Tube #1 (deemed to contain two distinct fibre-populations), the estimated median diameter for population 1 (p1) was 4.97 μm, and population 2 (p2) was 5.19μm (mean diameter: p1/p2 = 4.96/5.09 μm). Tube #4 was deemed to contain two blocks, each with a single fibre-population, but with distinct orientations. The median pore diameter in the first block (#4a) was 4.98 μm, and in the second block (#4b) was 5.04 μm (mean diameter: 4a/4b = 5.00/5.05 μm). Tube #5 was deemed to contain a single-fibre-population model, with a median diameter of 4.67 μm (mean diameter = 4.77 μm). Finally, Tube #6 was also deemed to contain two distinct fibre populations (median diameter: p1/p2 = 4.84/4.72 μm; mean diameter: p1/p2 = 4.89/4.76 μm). The fittings and parameter estimates were homogeneous across most voxels within the ROI (Figure 4).

**Figure 4.**
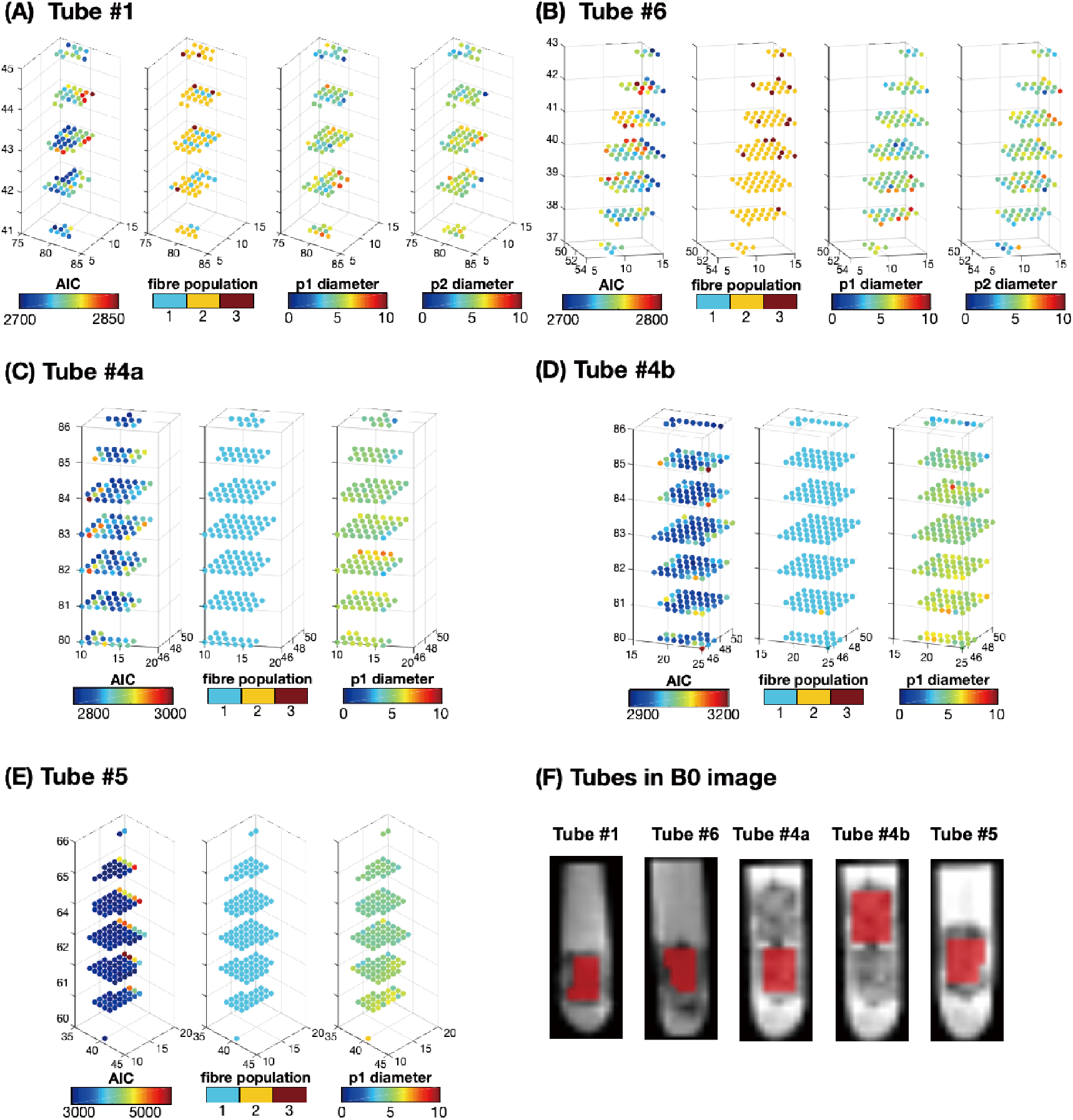
Fitting quality of fibre diameter estimations in five phantoms. To ensure fitting quality across different voxels, we examined 3 parameters including the Akaike Information Criterion (AIC), the number of distinct fibre populations (‘fibre population’), and the pore size diameter (‘p1 diameter’). This figure shows the manually-selected ROI in the 2D image slice, and the fitting results in the 3D scatter plots in a voxel-wise manner. Panels (A-B) are samples containing 2 distinct fibre populations, whereas panel (C-D) represents samples with a single fibre population.

#### Comparing the dMRI-derived estimates with SEM-derived estimates

The authors responsible for manufacturing the phantom (FZ, GP) confirmed that two of the six tubes contained orientationally-disperse samples (Tube #2 and #3), two contained samples with two fibre populations with crossing angles (Tubes #1 and #6), two tubes contained single fibre population blocks (Tubes #4 and #5), and one contained purely isotropic media (water). Figure 3A shows that the estimated orientations of the fibre populations were consistent with the ground truth fibre configuration; the median of crossing angles of Tube #1 and Tube #6 were 50.37° and 87.36° (Figure 3C), respectively, relative to the ground truth values of 45° and 90° (see Discussion regarding the precision of SEM/manufacturing). Figure 5 shows the Poisson fitting for all voxels within each ROI to recover the median diameter inside the fibre phantom across each scan. As noted above, due to the manufacturing process, the samples in Tube #1 and #6, and Tube #4a and #4b were derived from the same substrate, and thus there was only one set of SEM images available for each pair. For Tube #1 and #6, Tube #4a and Tube #4b, and Tube #5, the median (mean) diameters derived from SEM were 6.01 (6.43), 5.13 (5.82), and 5.38 (5.92) μm, respectively (Table 2). The dMRI-derived restricted signal fractions are similar to the values derived from SEM (total area of pores with diameter ≤ 15 μm / total area of pores with diameter in the range [0, ∞]), in which Tube #1 and Tube #6, #4, and #5 are 0.36, 0.47, and 0.57 respectively (Shown in Table 2).

**Figure 5.**
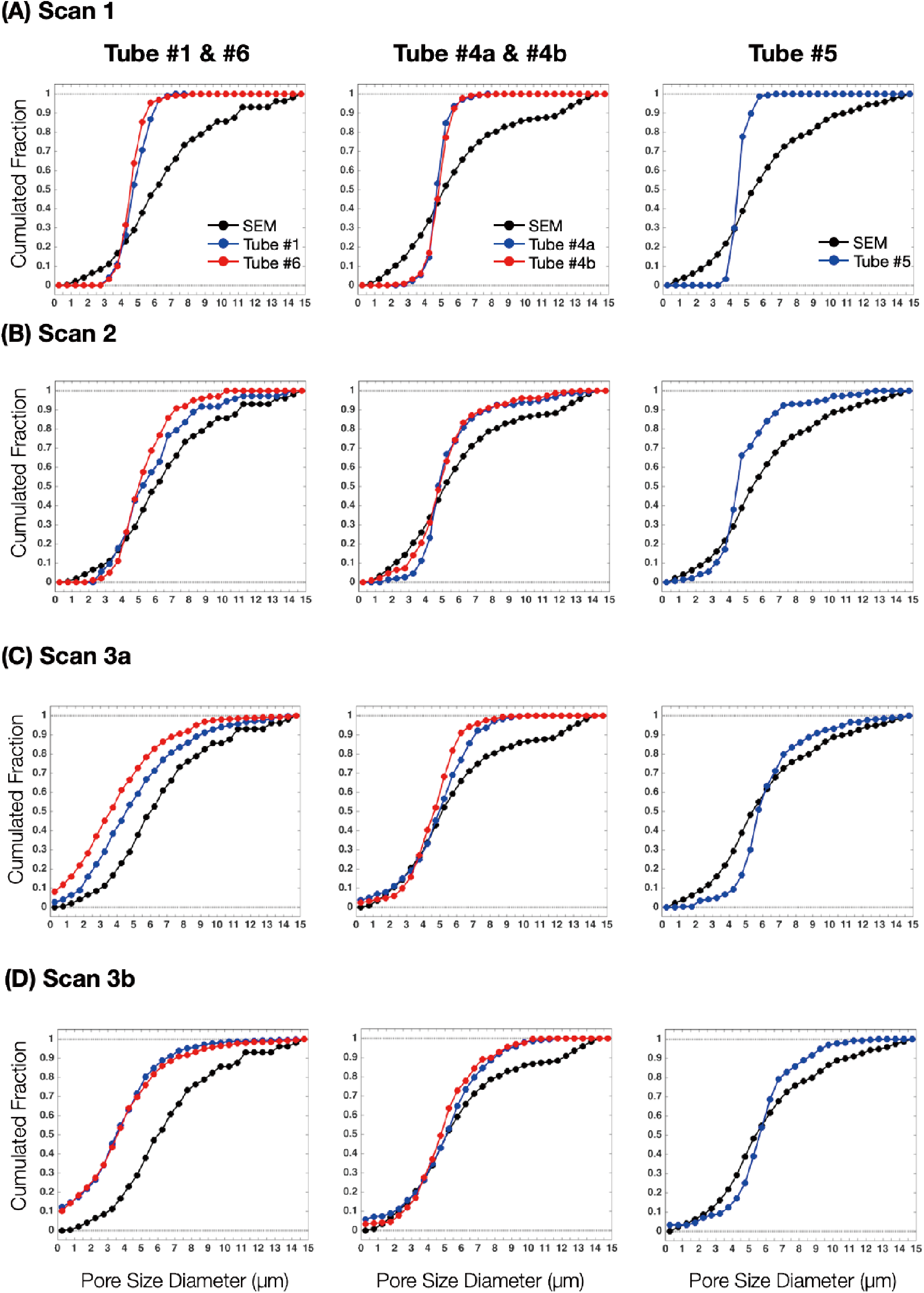
Cumulated fractions of the estimated fibre diameters from MRI and SEM. This figure shows the Poisson fitting for all voxels within each ROI for the different scans. The ground truth distribution of the pore size (estimated by SEM) is shown by the black line in each panel. Rows A to D represent the result from the different scans. The left column shows the two-fibre-population of Tube #1 and Tube #6, the middle column shows the single-fibre-population of Tube #4a and #4b, and the right column shows the single fibre-population of Tube #5.

The KS-test indicated that the distributions recovered from the first dMRI scan and SEM were significantly different (*P* < 0.01). No significant differences were found between dMRI and SEM results in Scan 2, Scan 3a, and Scan 3b, except for Tube # 4a in Scan 3b (Figure 5 and Table 3). The phantom characterization in different tubes is presented by the SEM micrographs (see Figure 2A-C) in which we observed that the phantom does contain some ‘extra-fibre’-like spaces. The histogram of pore size diameter against the area-weighted fraction was also shown in Figure 2D to depict the distribution of pore sizes in different phantom blocks.

**Table 3.**
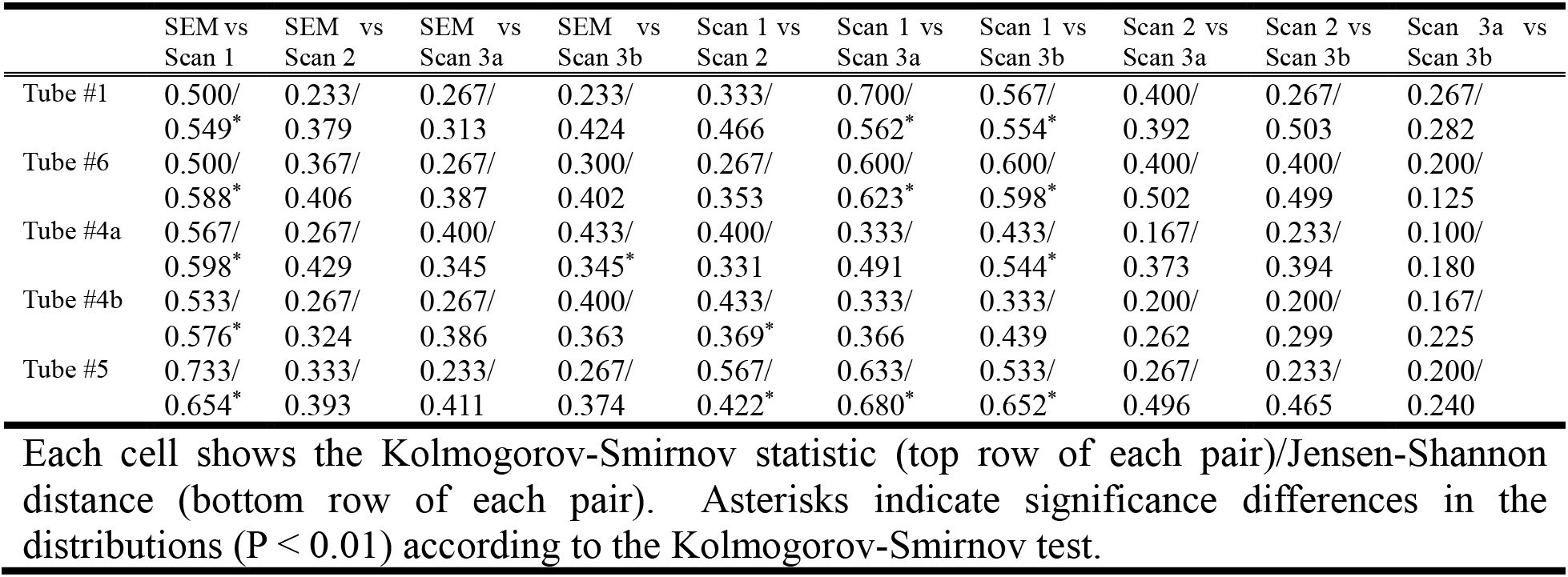
Equality between distribution of pore-size estimates obtained from SEM and different MRI scans evaluated by Kolmogorov-Smirnov test and Jensen-Shannon distance.

|  | SEM vs<br>Scan 1 | SEM vs<br>Scan 2 | SEM vs<br>Scan 3a | SEM vs<br>Scan 3b | Scan 1 vs<br>Scan 2 | Scan 1 vs<br>Scan 3a | Scan 1 vs<br>Scan 3b | Scan 2 vs<br>Scan 3a | Scan 2 vs<br>Scan 3b | Scan 3a vs<br>Scan 3b |
| --- | --- | --- | --- | --- | --- | --- | --- | --- | --- | --- |
| Tube #1 | 0.500/<br>0.549* | 0.233/<br>0.379 | 0.267/<br>0.313 | 0.233/<br>0.424 | 0.333/<br>0.466 | 0.700/<br>0.562* | 0.567/<br>0.554* | 0.400/<br>0.392 | 0.267/<br>0.503 | 0.267/<br>0.282 |
| Tube #6 | 0.500/<br>0.588* | 0.367/<br>0.406 | 0.267/<br>0.387 | 0.300/<br>0.402 | 0.267/<br>0.353 | 0.600/<br>0.623* | 0.600/<br>0.598* | 0.400/<br>0.502 | 0.400/<br>0.499 | 0.200/<br>0.125 |
| Tube #4a | 0.567/<br>0.598* | 0.267/<br>0.429 | 0.400/<br>0.345 | 0.433/<br>0.345* | 0.400/<br>0.331 | 0.333/<br>0.491 | 0.433/<br>0.544* | 0.167/<br>0.373 | 0.233/<br>0.394 | 0.100/<br>0.180 |
| Tube #4b | 0.533/<br>0.576* | 0.267/<br>0.324 | 0.267/<br>0.386 | 0.400/<br>0.363 | 0.433/<br>0.369* | 0.333/<br>0.366 | 0.333/<br>0.439 | 0.200/<br>0.262 | 0.200/<br>0.299 | 0.167/<br>0.225 |
| Tube #5 | 0.733/<br>0.654* | 0.333/<br>0.393 | 0.233/<br>0.411 | 0.267/<br>0.374 | 0.567/<br>0.422* | 0.633/<br>0.680* | 0.533/<br>0.652* | 0.267/<br>0.496 | 0.233/<br>0.465 | 0.200/<br>0.240 |

Each cell shows the Kolmogorov-Smirnov statistic (top row of each pair)/Jensen-Shannon distance (bottom row of each pair). Asterisks indicate significance differences in the distributions ( $P < 0.01$ ) according to the Kolmogorov-Smirnov test.

#### Repeatability of the dMRI-derived estimates

Table 3 shows the KS-test applied to different scans among different tubes for evaluating the long-term and short-term repeatability. The distribution obtained from Scan 1 is significantly different to those obtained from later scans, where all recovered distributions are broadly similar. The experimental design of the 3^rd^ scan session (i.e., no phantom re-positioning between the two scans) allowed estimates of short-term repeatability on a voxel-wise basis (Figure 6 and Table 4). For Tube #4a and Tube #4b, the voxel-level RC values of the estimated diameters are 4.13 μm and 4.10 μm, the ICC values are 0.519 and 0.361 respectively. The tubes containing samples with crossing fibre architectures (Tubes #1 and Tube #6) showed higher RC (Tubes #1 / Tube #6: 5.52 μm / 4.43 μm) and lower ICC values between repeat scans (ICC Tube #1 / Tube #6: 0.150 / 0.131). The sample-level RC value is 1.13 μm, the ICC value is 0.727, and the CV value is 11.3%.

**Table 4.** Repeatability between different MRI scans 3a and scan 3b.

| | RC ($\mu\text{m}$) | ICC | CV(%) |
| --- | --- | --- | --- |
| **Voxel-level repeatability** |  |  |  |
| Tube #1 | 5.52 | 0.150 | 58.9 |
| Tube #6 | 4.43 | 0.131 | 52.6 |
| Tube #4a | 4.13 | 0.519 | 40.6 |
| Tube #4b | 4.10 | 0.361 | 42.2 |
| Tube #5 | 4.75 | 0.201 | 38.8 |
| **Sample-level**<br>**repeatability** | 1.13 | 0.727 | 11.3 |

RC = repeatability coefficient; ICC = intra-class correlation coefficient; CV = within-voxel coefficient of variance; Asterisks are shown to indicate that $P < 0.01$

**Figure 6.**
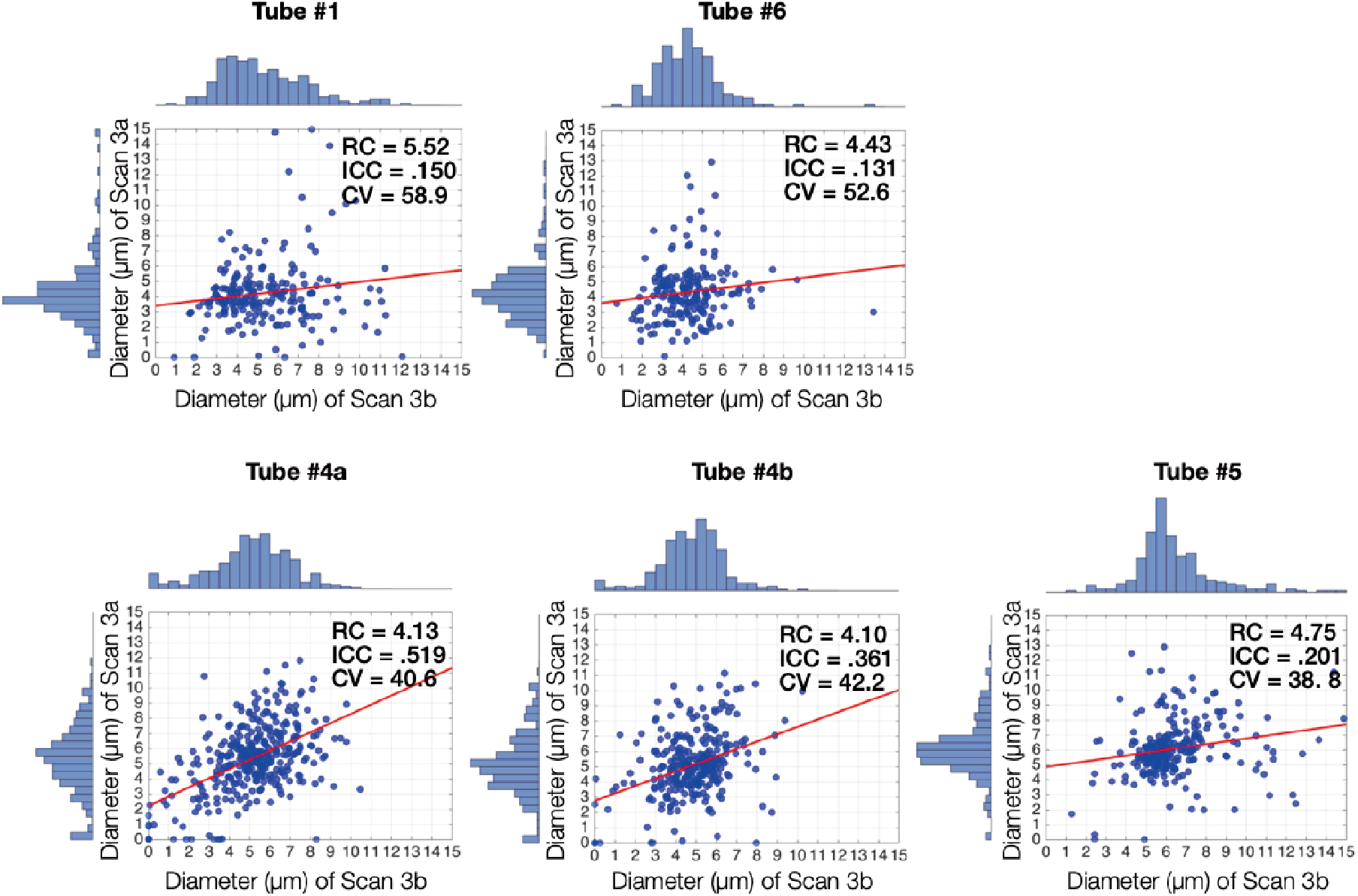
The intra-class correlation coefficient between immediate scan and rescan. The upper row plots show the estimated diameter from scan 3a against that from scan 3b of block samples with crossing orientation (in Tube #1 and #6), whereas the bottom row plots are block samples with single orientation (in Tube #4 and #5). The line of best fit between the data from the repeated scans is shown in red. RC: repeatability coefficient (μm); ICC: intra-class correlation coefficient; CV: within-voxel coefficient of variance (%).

## Discussion

This single-blind study used a 3T Connectom human MRI scanner, advanced modelling, and a co-electrospun hollow PCL-PSi microfibre phantom to establish the reliability of microfibre diameter estimates in a scan time < 1hr.

The results demonstrate fibre orientation and median pore size estimates which are highly comparable with results obtained by SEM, demonstrating the validity and robustness of the microstructural imaging pipeline with the phantom configuration used here. Compared with others in the literature, this phantom confers several advantages. The inner diameter approximates the median of the <u>range</u> of diameters within the human white matter (0.25~10 μm)^48^ (although please see ‘Limitations’ below). Moreover, the pore shape is more comparable to that seen *in vivo* making the phantom more ‘biomimetic’ than other phantoms developed to date. Finally, (see Figure 2), the substrate contains larger extra-fibre ‘voids’ between the restricting geometries (a result of the manufacturing process) where the diffusion path-length will be considerably longer than for spins trapped within the intra-fibre pores. Thus, the phantom has surrogate ‘extra-fibre’ compartments as well as intra-fibre compartments, which again pushes the properties closer to that of real tissue.

A previous study used a phantom comprising both extra- and intra-fibre compartments with a uniform inner diameter (12 ± 0.9 μm)^10^. The same group recently constructed a phantom with an inner diameter of 0.8 μm^14^, and have started to fashion cross-fibre configurations. However, no quantitative estimates of pore-size in these more complex configurations have been reported. Validation of such estimates in phantoms with complex architectures is necessary since human white matter fibre bundles are not perfectly co-aligned, even in ‘single fibre’ populations, and 60 ~ 90% of voxels contain multiple fibre orientations^42^. Thus, the acquired signal in each voxel may originate from the restricted water across several fibre bundles in different orientations or even more complicated geometrical configurations (e.g., axonal diameter), increasing uncertainty in fibre orientation estimates^49^. Such partial volume effects confound the estimation of fractional anisotropy and fibre diameter in restricted volumes.

The present study extends previous work by validating the AxCaliber3D framework using a more sophisticated phantom that includes a range of pore sizes (with a median around 5 μm), and both single and crossing fibre-orientations (about 45° and 90°). Regarding non-crossing single fibre conditions, the AxCaliber framework has been shown to recover axonal diameter distributions accurately^7,50^. In the current study, to account for crossing fibre configurations, we utilized AxCaliber3D^31^ with a continuous Poisson pore size distribution^30^ to resolve diameters. On the whole, regardless of whether the sample contained one or two fibre populations, the recovered fibre orientation and pore-size estimates agreed well with measurements obtained by direct SEM.

Good repeatability is critical to quantitative MRI research to provide stable metrics that are less influenced by measurement instability. Long-term repeatability facilitates the study of subtle longitudinal changes in pore size, while short- (and long-) term variability both impact the random errors and precision of the estimation model. Our results showed inconsistency of estimated pore-size distributions between the first scan session and other scan sessions. Across the long-term scans, it was not possible to perform a voxel-by-voxel comparison due to the difference in the phantom positioning. Further, images with lower SNR may introduce uncertainties in orientation and restricted-diffusion signal fraction estimation and thus result in variations in pore-size distribution^51^. That is, differences in estimated pore-size distribution might be explained, in part, by differences in the SNR.

We observed a reduction in SNR of approximately 25% over one year (between scans 2 and 3), although no difference was observed between scans 1 and 2. The source of this variation in SNR is unclear but could possibly reflect changes in the phantom material. For example, in the co-electrospinning manufacturing process, the core solution is PEO in water while the shell solution is PCL in Chcl3+DMF. The hollow microfibres are formed *in situ* after the evaporation of the solvents in both the shell and core. The PEO polymer is assumed to deposit on the inner surface of the resultant hollow PCL fibres but is not removed before the phantom is assembled. It can be expected that PEO would dissolve gradually in water, when the hollow fibres are filled with water. However, PEO has a very high molecular weight (900 kg/mol) and dissolves very slowly at room temperature. The PCL polymer in the microfibre shell is subject to hydrolytic degradation, taking 2-4 years for a complete degradation, depending on the initial molecular weight and surrounding fluid^52^. Therefore, considering the one-year gap between the second and third scan sessions, a certain level degradation of PCL polymer can be also expected in the water-filled phantom. This may have shortened T_2_, leading to a reduction in SNR and is worthy of further investigation, but is beyond the scope of the current work.

Duval, et al.,^53^ previously demonstrated stable AxCaliber estimates in the spinal cord of healthy human participants, with correlation coefficient (r) 0.64. We note that in the spinal cord, the axons tend to be largely co-axial. In our study, pore-size estimates demonstrated good repeatability at the sample-level (ICC = 0.727, RC = 1.13 μm), whereas the repeatability of pore-size estimates at the voxel-level was considerably better for ‘single orientation’ samples than for those containing multiple fibre orientations. However, the short-term repeatability remained poor in both cases, which suggests the uncertainty at the voxel level that may still be affected by the errors from many parameters fitting in the model or potential residual misalignment in the scan-rescan test. Nevertheless, pore size and fibre orientations were estimated accurately by the proposed framework, but the repeatability at the voxel level should be re-examined and improved in the future.

### Limitations of the Study and Future Directions

While our work demonstrated the strength and reliability of AxCaliber3D model to resolve the complex fibre architectures <u>in this particular substrate</u>, it is important to keep in mind that this study also has several limitations. Most importantly, we caution against full extrapolation to ‘axon diameter’ mapping. While this phantom is a definite move *towards* the white matter properties, it would be premature (and incorrect) to conclude that this work fully licenses claims about the validity of AxCaliber3D for estimating axon diameters in all of white matter.

The most obvious hurdle preventing these claims is that the pore sizes remain considerably larger than the *<u>modal</u> ex vivo* white matter axons in the human brain. Further work is needed to manufacture pores with a smaller internal diameter. Secondly, there was no explicit attempt to control the temperature of the scan room during data acquisition. Lack of temperature control may have led to differences in the diffusivities of the phantom substrate between scans which may have affected the precision/ repeatability of the microstructural estimates. Although we did not anticipate this to be a major contributor, future work should record the real-time temperature during scanning to clarify such a possible confound. Moreover, we observed changes in the diffusion MRI characteristics of the phantom materials between scans 1 and 2 that are challenging to explain. We considered degradation of the phantom, but the only potential evidence of this is a change of around 25% in SNR that was observed between scans 2 and 3, over a period of 1 year; this does not explain the differences between scans 1 and 2. Additionally, as can be seen in Figure 5 and Table 3, the cumulative histograms of fibre diameters for the SEM measurements of the phantom (measured before any of the MRI scans) match the cumulative histograms of fibre diameters from the diffusion MRI scans at time points 2, 3a, and 3b well. The diffusion MRI cumulative histograms from scans 2, 3a, and 3b also match each other well but do not match the diffusion MRI histograms from scan 1. The cause of this outlier behaviour for the diffusion measurements at scan 1 requires further investigation in future longitudinal studies; at this time, we are unable to rule out potential short-term issues with water penetration into the phantom material, scanning temperature variations, or errors in the diffusion MRI data acquisition as potential causes.

Thirdly, the phantom was not explicitly designed to mimic the relative size and shape of the extra-axonal space seen in tissue, and thus estimates of tortuosity and extra-axonal time-dependence are unlikely to reflect the situation *in vivo^12,30,54^*. Fourth, the degree to which water exchanges across the fibre membranes is currently unknown, although a reasonable degree of restriction is apparent in the clear presence of time-dependent diffusion^28^. Finally, due to the way in which the phantom is manufactured, the substrate is heterogeneous and thus the control of the ‘ground truth’ fibre angle and pore size is imprecise, which adds uncertainty to the cross-validation process. Thus, achieving perfect agreement between the dimensions extracted from the SEM of a subsample of the material and the sample used for imaging can be challenging.

Despite these limitations, this study can be considered as a useful step in the evolution of validating pore size estimates in complex geometries on a human MRI scanner. Similar validations in non-uniform pore phantoms in relatively short scanning times will enlighten the clinical practice of microstructural imaging in different living tissues including, but not limited to, the prostate^55^ and muscle fibres^56,57^.

Owing to the blind nature of the experiment, the sample geometry and restricted diffusivity were totally unknown *a priori*. Thus, a wide range of diffusion times (diffusion time: 19 ~ 90 ms) was used to maximize the sensitivity of diffusion displacement to possible pore sizes^58^. If more information is known *a priori*, the acquisition protocol could be optimised accordingly, including a reduction in total acquisition time. This is likely to lead to improved precision of microstructural parameter estimation and may therefore also improve repeatability.

In summary, by spanning multiple diffusion times and gradient strengths on an ultra-strong gradient scanner, we successfully estimated the fibre architectures that had expected pore sizes lower than 10 μm (around 5 μm) in both single-aligned fibre populations and in populations of crossing fibres within a new biomimetic phantom with non-uniform cross-sections, which more closely mimics the white matter features than previously-employed simple geometric phantoms. Our microstructure measurements show good agreement with the new generation diffusion phantom and support validity for microstructure quantification of complex environment at the micron level. Future work is underway to validate pore-size estimates in phantoms with more crossing populations (including completely random), variable ‘extra-cellular’ volume fractions and smaller non-uniform pore-size diameter than studied here.

## Acknowledgements

The data were acquired at the UK *National Facility for In Vivo MR Imaging of Human Tissue Microstructure* funded by the EPSRC (grant EP/M029778/1), and The Wolfson Foundation. DKJ and MD were both supported by a Wellcome Trust Investigator Award (096646/Z/11/Z) and a Wellcome Trust Strategic Award (104943/Z/14/Z). We thank Dr John Evans for his help to acquire the repeat-scan MRI data. We thank Dr Robbert Harms for his assistance with embedding the pore-size estimation routine into the MDT toolbox. FLZ was supported by NIHR UCLH Biomedical Research Centre (BRC) grant and UK-MRC ImagingBioPro grant (MR/R025673/1). GJM Parker has a shareholding and part time appointment and directorship at Bioxydyn Ltd. which provides MRI services. GJM Parker also has a shareholding and directorship at Queen Square Analytics Ltd. which provides MRI services. The authors gratefully acknowledge the assistance from Prof. Julie E. Gough and Prof. Gareth R. Williams with access to lab space and electron microscopy facilities at The University of Manchester and University College London, respectively. CPL was supported by the Ministry of Science and Technology (MOST) of Taiwan (grants MOST 109-2634-F-010-001, MOST 108-2321-B-010-013-MY2, MOST 108-2321-B-010-010-MY2, and MOST 108-2420-H-010-001). Author H (Chu-Chung Huang), C-CHH (Chih-Chin Heather Hsu) and FLZ (Feng-Lei Zhou) contributed equally to this work.

