## Supplemental Figure S1 for "Validating Pore Size Estimates in a Complex Microfibre Environment on a Human MRI System"

### SUPPLEMENTARY MATERIAL

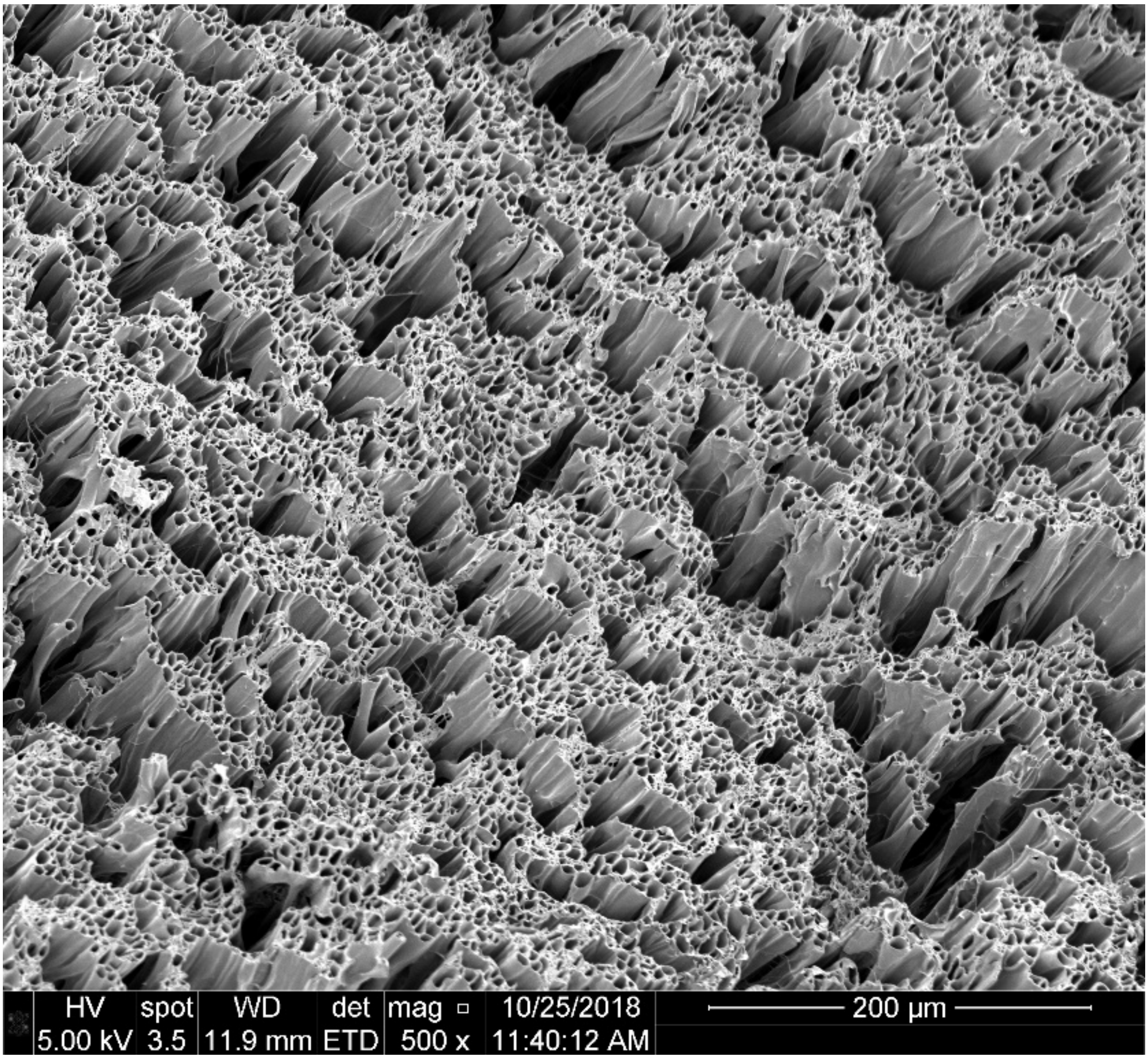


**Figure S1.** Larger field of view SEM image showing the irregularity of the manufacturing process, including the large ‘extra-fibre’ pores
